# Inhibition of myeloperoxidase attenuates thoracic aortic aneurysm formation in Marfan disease

**DOI:** 10.1101/2022.11.24.517172

**Authors:** Dennis Mehrkens, Felix Sebastian Nettersheim, Felix Ballmann, Jil Bastigkeit, Alexander Brückner, Johannes Dohr, Simon Geissen, Lauren De Vore, Patrik Schelemei, Felix Ruben Picard, Malte Kochen, Simon Braumann, Wiebke Kreuzberg, Alexander Hof, Henning Guthoff, Adrian Brandtner, Benedicta Quaye Mensah, Maarten Groenink, Mitzi van Andel, Arnout Mieremet, Susanne Pfeiler, Norbert Gerdes, Ulrich Flögel, Laura-Maria Zimmermann, Gerhard Sengle, Marie-Lisa Eich, Birgid Schömig-Mariefka, Matti Adam, Bernd K Fleischmann, Daniela Wenzel, Vivian de Waard, Anna Klinke, Stephan Baldus, Martin Mollenhauer, Holger Winkels

## Abstract

Marfan syndrome (MFS) is the most prevalent inherited connective tissue disorder, still remains uncurable, and is characterized by high mortality at early age driven by dissection and rupture of thoracic aortic aneurysms. MFS is caused by mutations in the fibrillin-1 gene and aberrant TGFß signaling.

Here we addressed whether myeloperoxidase (MPO), a leukocyte derived enzyme with potent matrix modulating properties also influences the aortic phenotype in MFS.

MFS patients displayed increased circulating MPO levels compared to controls as well as marked aortic MPO deposition. In an MFS mouse model, MPO induced inflammatory endothelial activation and endothelial to mesenchymal transition which triggered aortic leukocyte recruitment. Moreover, MPO directly contributed to adverse extracellular matrix remodeling by promoting oxidative stress and nitration of proteins within the vascular wall. Genetic MPO deficiency and pharmacological MPO inhibition attenuated MFS-related aneurysm formation. We herein identify MPO as a critical mediator of MFS-related thoracic aortic aneurysm formation and - in the absence of any pharmacological treatment so far in this disease - a first anti-inflammatory target to modulate disease progression.

## Introduction

Marfan syndrome (MFS) is the prototype of hereditary autosomal dominant connective tissue disorder with a prevalence of 1:5000.^1^ Its prognosis is driven by the vascular phenotype. Due to thoracic aortic aneurysm (TAA) formation and subsequent dissections or ruptures of the aorta, MFS is associated with premature mortality.^1^ MFS is caused by mutations in the fibrillin-1 gene (*Fbn1*) encoding the extracellular matrix (ECM) protein fibrillin-1.^2^ Fibrillins are the major components of microfibrils, complex extracellular structures that provide the core scaffold for amorphous elastin in elastic fibers. In MFS, *Fbn1* deficiency or mutations cause elastin strand breaks, which crucially contribute to TAA formation and dissection.^3^ Several mechanisms underlie elastic fiber fragmentation. Nitric oxide (NO) signaling is disturbed in MFS and sufficed to cause extracellular fibrillin fragmentation.^4^ Mutations in fibrillin-1 adversely affect the structural integrity of fibrillin microfibrils and induce degradation by matrix metalloproteinases (MMP). In vitro, fibrillin-1 fragments stimulate expression of MMPs.^5,6^ The degradation of microfibrils contributes to progressive tissue destruction.^7^ Besides their effect on structural properties, microfibrils play a critical role in tissue homeostasis and aid in sequestering extracellular growth factors and cytokines, including transforming growth factor beta (TGF-β), thereby preventing aberrant activity.^8^ TGF-β is an important mediator of vascular homoeostasis with in part dichotomous effects in aortic pathology: On the one hand it prevents inflammatory abdominal aortic aneurysm formation^9^ and early aortic disease development in MFS.^10^ Complete abrogation of TGF-β signaling aggravates MFS-related aortic dilatation.^11^ On the other hand, TGF-β overactivation promotes MFS-related TAA formation at later disease stages.^10,12,13^ Several mechanisms modulating TGF-β levels may influence disease severity. Plasma from MFS patients shows signs of oxidative stress^14^ and increased levels of reactive oxygen species (ROS) in human Marfan aneurysms induce TGF-β release.^15^ Additionally, ROS target cytoskeletal proteins, e.g. smooth muscle α-actin (ASMA), a key component controlling cell adhesion and contractility in vascular smooth muscle cells (VSMC). Augmented ROS levels in the aortic wall might thus contribute to the pathogenesis of MFS.^16^ Several inflammatory processes can lead to elevated ROS, MMP, and TGF-ß bioavailability. Recombinant fibrillin-1 fragments and aortic extracts from MFS mice stimulate macrophage chemotaxis,^17^ suggesting that leukocyte infiltration might drive the pathogenesis of MFS. Immunohistochemical analysis of aortic sections from MFS patients and mice confirmed leukocyte infiltration in the aortic tunica media and aorta, respectively.^17,18^ The role of the innate immune system and its mediators is - in striking contrast to many other types of cardiovascular diseases - in MFS poorly understood.

Myeloperoxidase (MPO) is one of the principal enzymes released upon activation of PMN, monocytes, and macrophages. MPO interacts with surface receptors expressed on endothelial cells (EC),^19^ which are involved in transcytosis resulting in the deposition of catalytically active MPO in the subendothelial space.^20^ MPO uses hydrogen peroxide (H_2_O_2_) as a co-substrate and oxidizes halides, in particular chloride, thereby generating hypochlorous acid (HOCl).^21^ HOCl has been linked to activation of Rho-kinase thereby impairing vasorelaxation and increasing vascular tone, respectively.^22^ Moreover, HOCl activates MMPs and MAP kinases in fibroblasts, which provokes a pro-synthetic phenotype.^23^ MPO further weakens stability of the endothelial glycocalyx and mediates its shedding.^24^ Due to its cationic charge and independent of its catalytical activity, MPO directly attracts PMN to the vascular wall and promotes proinflammatory leukocyte adhesion and extravasation into the vessel wall.^24^ Of note, inhibition of MPO has been shown to attenuate vascular pathologies in preclinical models including atherosclerosis^25^ and endothelial dysfunction.^26^ The role of MPO in MFS is unknown.

MFS patients are treated with ß1-receptor blockers or Angiotensin-II-receptor antagonists (ARBs). As of today, none of these therapies has been shown to critically alter the course of the disease in patients. Given the fatal course of the disease and the limited pharmacological treatment options, characterization of the contributing mechanisms including the innate immune system is important and may pave future avenues of treatment. In this study, we aim to unravel the role of MPO in the progression of MFS to identify potential novel treatment strategies for MFS patients.

## Material and Methods

We analyzed systemic MPO levels in patients with MFS and studied the pathomechanistic role of MPO in MFS in a mouse model. Heterozygous missense mutation *Fbn1^C1041G/+^* (MFS) mice were crossed with MPO-deficient mice (MFSx*Mpo*^-/-^) or treated with an oral MPO inhibitor. TAA formation was examined by ultrasound, MRI and histological analyses. Furthermore, the impact of MPO on MFS-related endothelial activation and leukocyte recruitment was analyzed by intravital microscopy and RNA sequencing of endothelial cells. The expanded methods are provided in the Supplemental Material.

## Results

### Myeloperoxidase is elevated in MFS patients and mice

To investigate the impact of MPO in MFS, we first measured MPO plasma levels, which were significantly increased in MFS patients compared to healthy controls (**Fig. 1A**). Concomitantly, the ascending aortic sections from MFS patients showed MPO deposition (**Fig. 1B**). Mice with the Marfan genotype (MFS) had elevated plasma and tissue immobilized MPO in the ascending aorta in comparison to wild-type controls (WT) (**Fig. 1C, D**). As expected, MPO was not detectable in MFSx*Mpo*^-/-^ mice. The elevated MPO levels in MFS mice coincided with increased numbers of circulating leukocytes and neutrophils (**Fig. 1E, F, Sup Fig. 1**).

**Figure 1.**
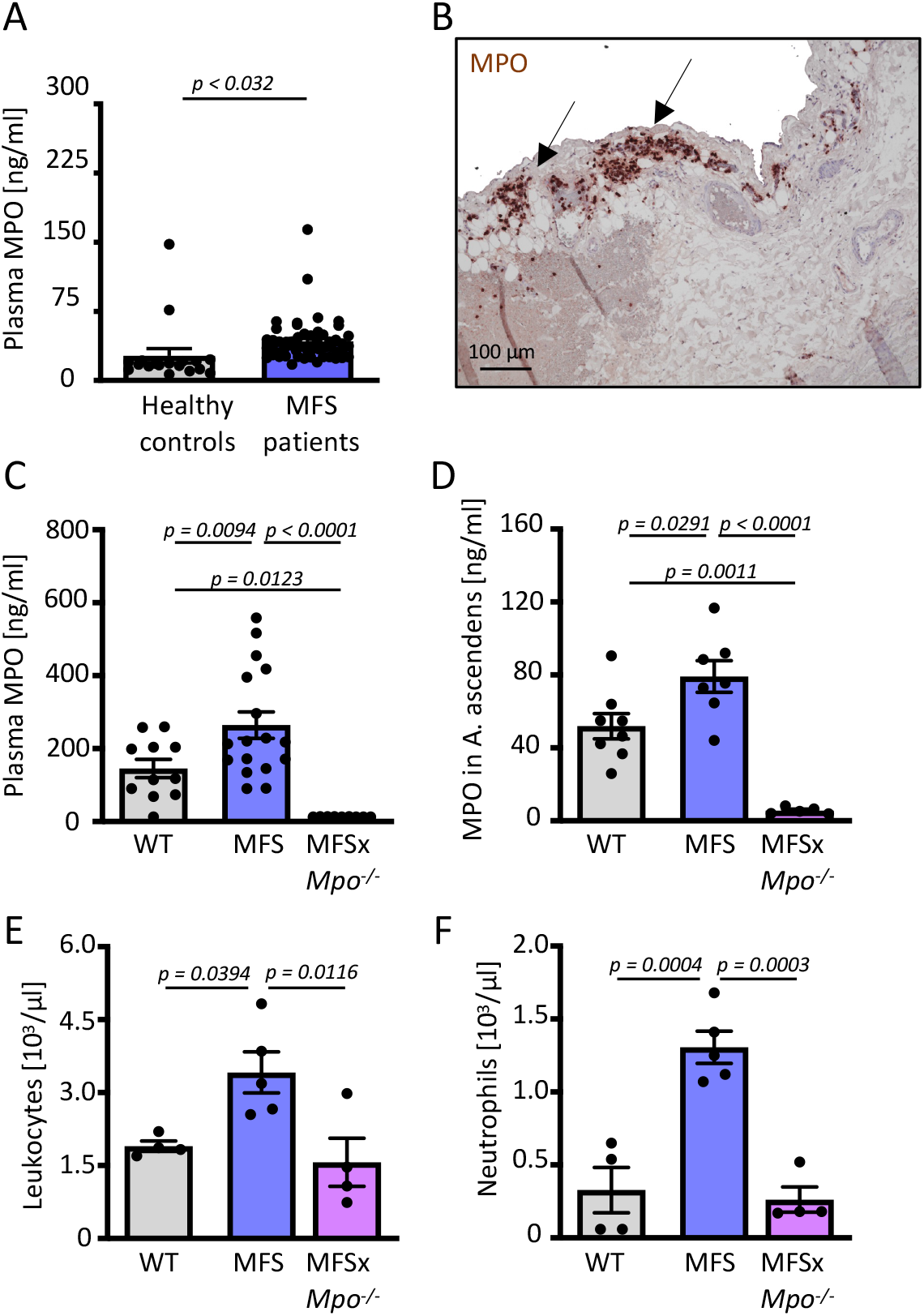
Myeloperoxidase is elevated in MFS patients and mice. (A) Plasma myeloperoxidase (MPO) levels in Marfan patients (MFS, N=53) and healthy controls (N=18). (B) Representative image of MPO reactivity in *aorta ascendence* cross sections from an MFS patient. Scale bar indicates 100 μm. (C) MPO in plasma (WT N=11, MFS N=17, MFSx*Mpo*^-/-^ N=9). (D) MPO levels in the thoracic aortic arch (WT N=8, MFS N=7, MFSx*Mpo*^-/-^ N=5). (E,F) Circulating white blood cell (E) and neutrophil counts (F) (WT N=4, MFS N=5, MFSx*Mpo*^-/-^ N=4). Data is presented as mean ± SEM. Statistical significance was determined by (A) the Mann-Whitney test or (C-F) by ordinary oneway ANOVA followed by Tukey’s multiple comparison test. Wildtype (WT), Fbn1^C1041G/+^ (MFS), MPO-deficient MFS (MFSx*Mpo*^-/-^).

### Genetic myeloperoxidase-deficiency attenuates aortic dilatation and arterial stiffness in MFS mice

We next sought to determine whether MPO is functionally relevant for MFS. Echocardiographic analyses of the aortic root and the ascending aorta confirmed aortic dilatation in 12-14-week-old MFS animals in comparison to WT controls (**Fig. 2A-E**). These findings were supported by magnetic resonance imaging (MRI) indicating an increased aortic annulus area (**Fig. 2F, G**) in MFS mice. Pulse wave propagation velocity, a measure of arterial stiffness and indicator of progressive aortic dilatation, was also elevated in MFS mice (**Fig. 2H, I**). MFSx*Mpo*^-/-^ displayed attenuated aortic dilation and arterial stiffness (**Fig. 2A-I**)

**Figure 2.**
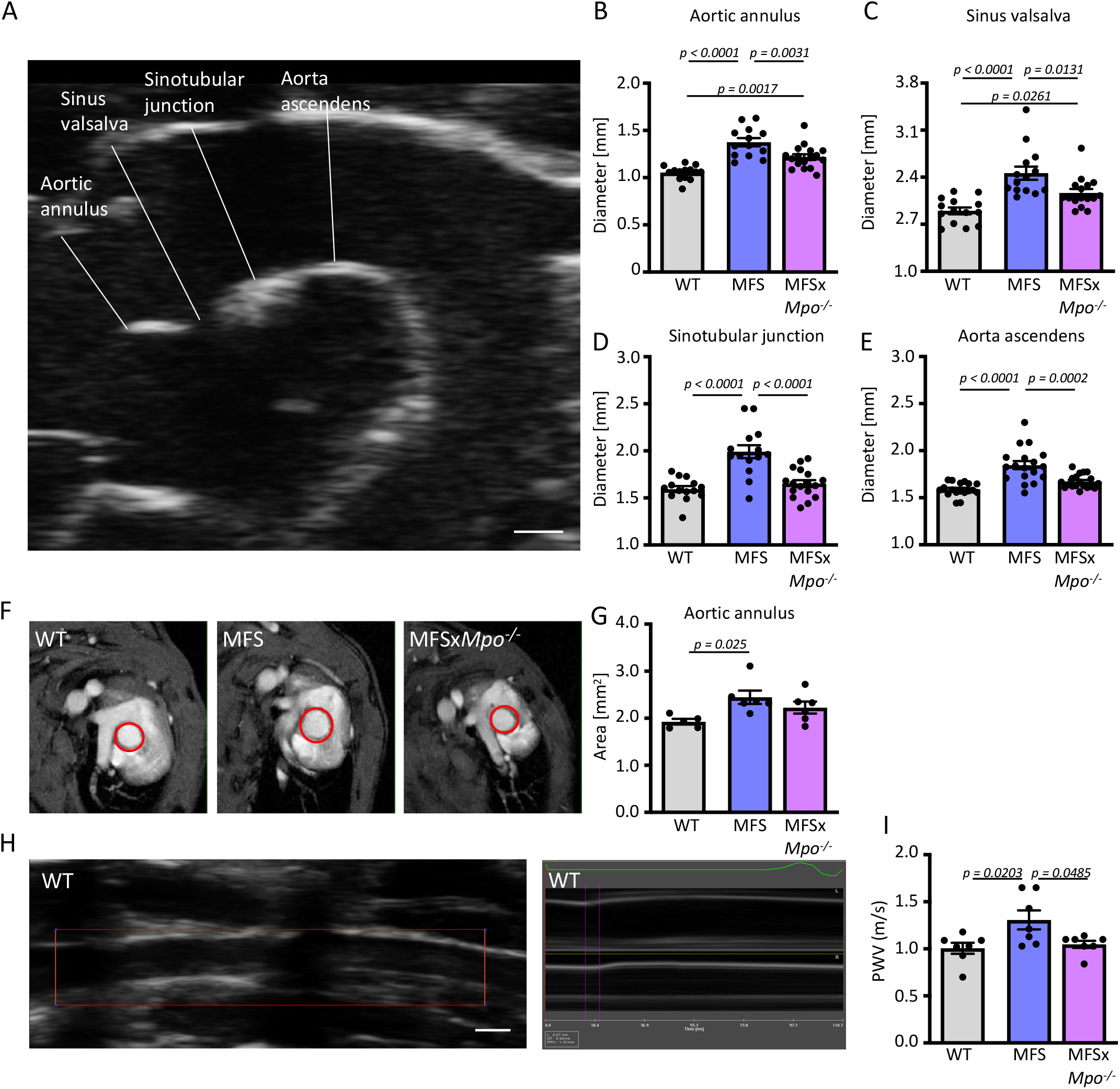
Genetic MPO deficiency attenuates MFS-related aortic dilation and stiffening. (A) Representative 2D echocardiographic image with diameter measurements at the position of the (B) aortic annulus, (C) sinus Valsalva, (D) sinotubular junction, and (E) aorta ascendens in 12-14-week-old WT, MFS, and MFSx*Mpo*^-/-^ mice. (WT N=14, MFS N=14, MFSx*Mpo*^-/-^, N=17). Scale bar indicates 1 mm. (F) Representative magnetic resonance imaging microphotographs and (G) quantification of the aortic annulus area WT N=5, MFS N=6, MFSx*Mpo*^-/-^, N=6). (H) Representative 2D echocardiographic image of WT aorta ascendens and (I) quantification of the pulse propagation velocity (WT N=7, MFS N=7, MFSx*Mpo*^-/-^, N=7). Scale bar indicates 500 μm. Data is presented as mean ± SEM. Statistical significance was determined by ordinary one-way ANOVA followed by Tukey’s multiple comparison test. Wildtype (WT), Fbn1^C1041G/+^ (MFS), MPO-deficient MFS (MFSx*Mpo*^-/-^).

### Myeloperoxidase-deficiency protects from aortic structural remodeling and oxidative stress in MFS mice

We next determined the effect of MPO deficiency on MFS-induced structural and inflammatory responses within the aortic wall. Aortic sections were histologically analyzed by Van-Giesson staining to determine elastin strand breaks as an indicator of disease progression. As expected, MFS mice showed increased fragmentation of elastic fibers in comparison to WT mice, while elastic fiber fragmentation in MFSx*Mpo*^-/-^ mice was comparable to WT level (**Fig. 3A**). Immunofluorescence staining of aortic sections showed strong 3-nitrotyrosine abundance - a potential product of MPO-derived nitrating species like nitric oxide^27^ and biomarker for oxidative stress - in MFS vs. WT and MFSx*Mpo*^-/-^ mice (**Fig. 3B**). An increase in the proteolytic activity of MMP2 and 9 has been linked to MPO-activity^28^ and extracellular matrix (ECM) degradation.^29^ In-gel and in-situ zymography revealed elevated proteolytic MMP2/9 activity in thoracic aortas of MFS mice, which was absent in WT and MFSx*Mpo*^-/-^ mice (**Fig. 3C, Sup Fig. 2**). ROS are further a mediator of cell death including apoptosis. Thoracic aortic sections from MFS in comparison to WT and MFSx*Mpo*^-/-^ mice harbored significantly more apoptotic cells as assayed by immunoreactivity for cleaved caspase 3 (**Fig. 3D**).

**Figure 3.**
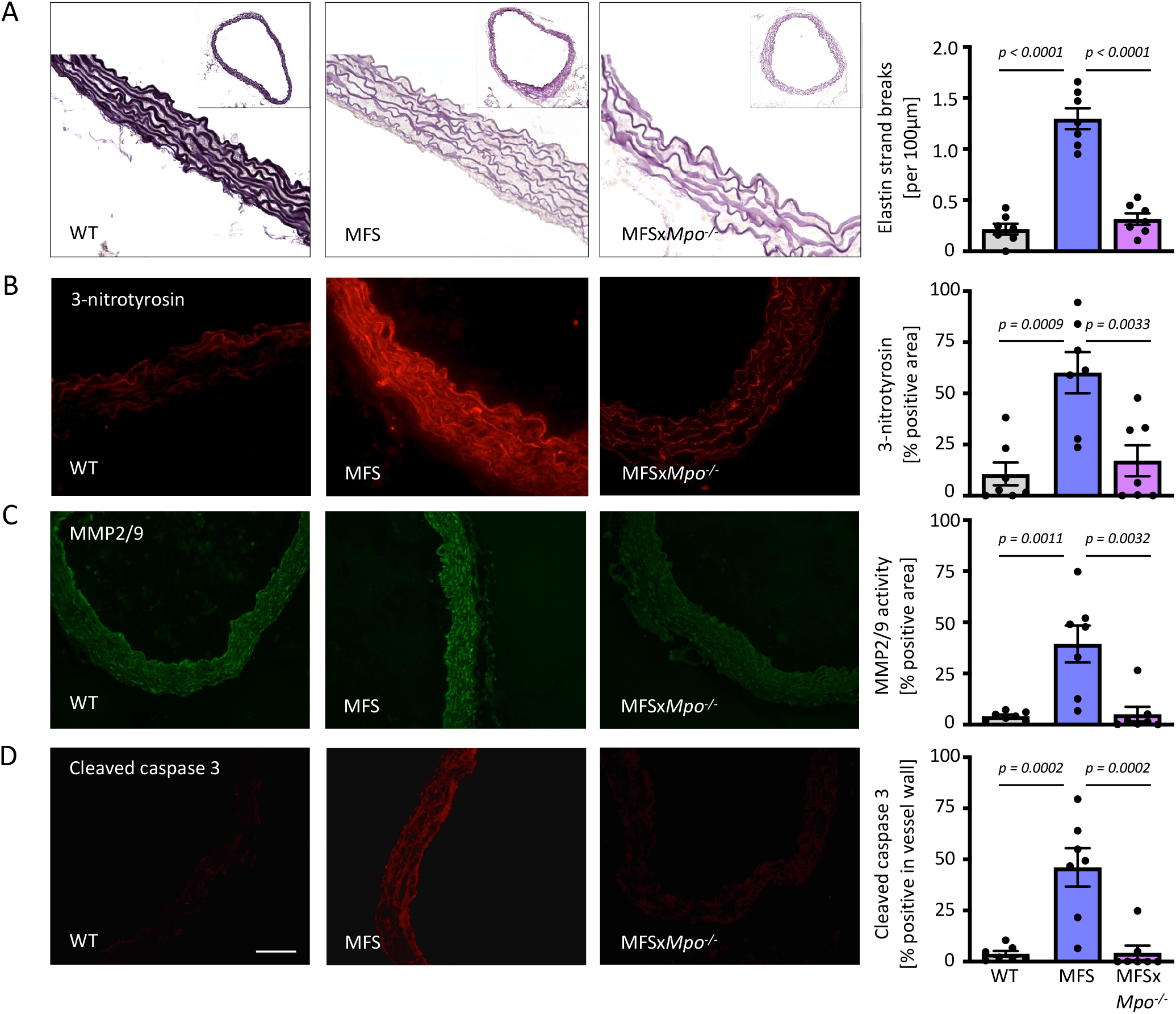
Genetic MPO deficiency attenuates aortic remodeling in MFS. (A) Representative images of elastica van Gieson stained ascending aortic sections and average number of elastin strand breaks per 100 μm of elastic lamina. (B) Representative images of 3-nitrotyrosine staining (red) and quantification as percentage of DCFHDA positive area (C). Representative in situ zymographies of MMP2/9 activity in ascending aortic sections and quantification of the gelatinolytic area (bright green) as percentage of aortic media. (D) Apoptotic cell death in the ascending aorta analyzed by percentage of active caspase 3 positive area (red). For each staining N=7/group. Data are expressed as mean ± SEM. Statistical significance was determined by one-way ANOVA with Tukey’s multiple comparisons test. Scale bar indicates 50μm. Wildtype (WT), Fbn1^C1041G/+^ (MFS), MPO-deficient MFS (MFSx*Mpo*^-/-^).

### MFS causes endothelial cell inflammation, which is ameliorated by MPO deficiency

Increased ROS production, MMP activity and cellular apoptosis are hallmarks of an inflammatory response. We next sought to investigate how MFS alters endothelial cell function and might thus contribute to TAA formation. We therefore performed in silico analysis of previously published aortic single-cell transcriptomes from WT and MFS mice.^30^ We identified two endothelial cell clusters expressing the hallmark genes *Pecam1, Cldn5 and Ctla2a*,^31^ which were present in both groups (**Sup Fig. 3A, B**). The larger cluster EC cl.1 was characterized by the expression of classic EC-associated genes including *Esam* and *Tie1. Esam* encodes for a cell adhesion molecule, which regulates angiogenesis, endothelial permeability, and leukocyte transmigration.^32^ Tie1 is a receptor for angiopoietin that regulates the survival of ECs and angiogenesis.^33^ The smaller cluster EC cl.2 was characterized by *Nts2, Ccl21a, Fabp4*, and *Lyve-1* expression (**Sup Fig. 4B, Sup Table 1**). The hyaluronan receptor *Lyve-1* is particularly expressed in lymphatic endothelium,^34^ which also secretes the chemokine CCL21.^35^ It is thus likely, that EC cl.2 contains lymphatic ECs. We next performed differential gene expression of EC cl.1 and EC cl.2 to understand the biological differences caused in MFS mice. We detected 228 differentially regulated genes in cluster EC cl.1 and 196 in cluster EC cl.2. Both EC populations shared 55 upregulated and 43 downregulated genes in MFS (**Sup Fig. 3C, D**). Commonly upregulated genes in MFS were indicative of endothelial to mesenchymal transition including *Acta2, Tagln, Myl9*, and *Itga8*, as well as multiple collagen-encoding genes (**Sup Table 1**). Simultaneously, both EC clusters expressed lower levels of the EC marker genes *Pecam1, Cldn5*, and *Cdh5* (**Sup Table 1**). Both EC populations further expressed multiple genes involved in cell migration and inflammation (EC cl.1: *Cxcl12, Cxcl2, Vcan*; EC cl.2: *Ccl4, Cxcl1, Ncam1*) (**Sup Table 1**). Gene ontology analysis revealed significant changes in biological functions of both cell clusters in MFS (**Sup Fig. 3E, F, Sup Table 2**). Both EC clusters showed enrichment for pathways predominantly involved in inflammation (*granulocyte chemotaxis, endothelial cell migration*) and extracellular matrix remodeling (*positive regulation of transforming growth factor beta receptor signaling pathways, wound healing, collagen fibril organization, extracellular matrix organization, regulation of collagen metabolic process*). Furthermore, multiple endothelial cell developmental and differentiation pathways (*endothelium development, regulation of endothelial cell proliferation, regulation of endothelial cell migration*), as well as angiogenesis pathways (*vasculogenesis, regulation of angiogenesis, sprouting angiogenesis*) were downregulated in both EC cluster. Nitric oxide biosynthesis is critically involved in vascular function, also this pathway was found to be inactivated in MFS ECs (**Sup Table 2**).

To further uncover how MPO alters EC function in MFS, we isolated ECs from the ascending thoracic aortas from 12-14-week-old WT, MFS and MFSx*Mpo*^-/-^ mice and subjected these cells to in depth transcriptional analysis. In comparison to WT mice, ECs from MFS and MFSx*Mpo*^-/-^ mice differentially expressed 908 and 795 genes of which 246 were jointly upregulated and 72 downregulated (**Fig. 4A-F, Sup Table 3-5**). We next subjected the mutually exclusive up- and downregulated genes to pathway analysis (**Fig. 4G, Sup Tab 6**). We also observed in bulk transcriptome analysis predominant upregulation of inflammatory pathways in MFS vs. WT EC including *inflammasome complex assembly, cellular response to type I interferon*, and *C-C chemokine receptor activity*. Other predominant pathways comprised cell cycle terms including *establishment of spindle localization* and *negative regulation of mitotic cell cycle phase transition*. In contrast, these pathways were not upregulated in ECs from MFSx*Mpo*^-/-^ mice, which were transcriptomically enriched for developmental pathways (*neural tube development, nephron development*). We also found that ECs from MFS and MFSx*Mpo*^-/-^ mice were both altered in their metabolism and respiration, as they showed downregulated respiration (*cellular respiration, aerobic respiration, oxidative phosphorylation, electron transport chain*) and reduced metabolic pathways (*pentose-phosphate shunt, triglyceride metabolic process, glucose metabolic process, nucleotide metabolic process*), as previously observed for MFS mouse aortas and human MFS aortic smooth muscle cells.^36,37^

**Figure 4.**
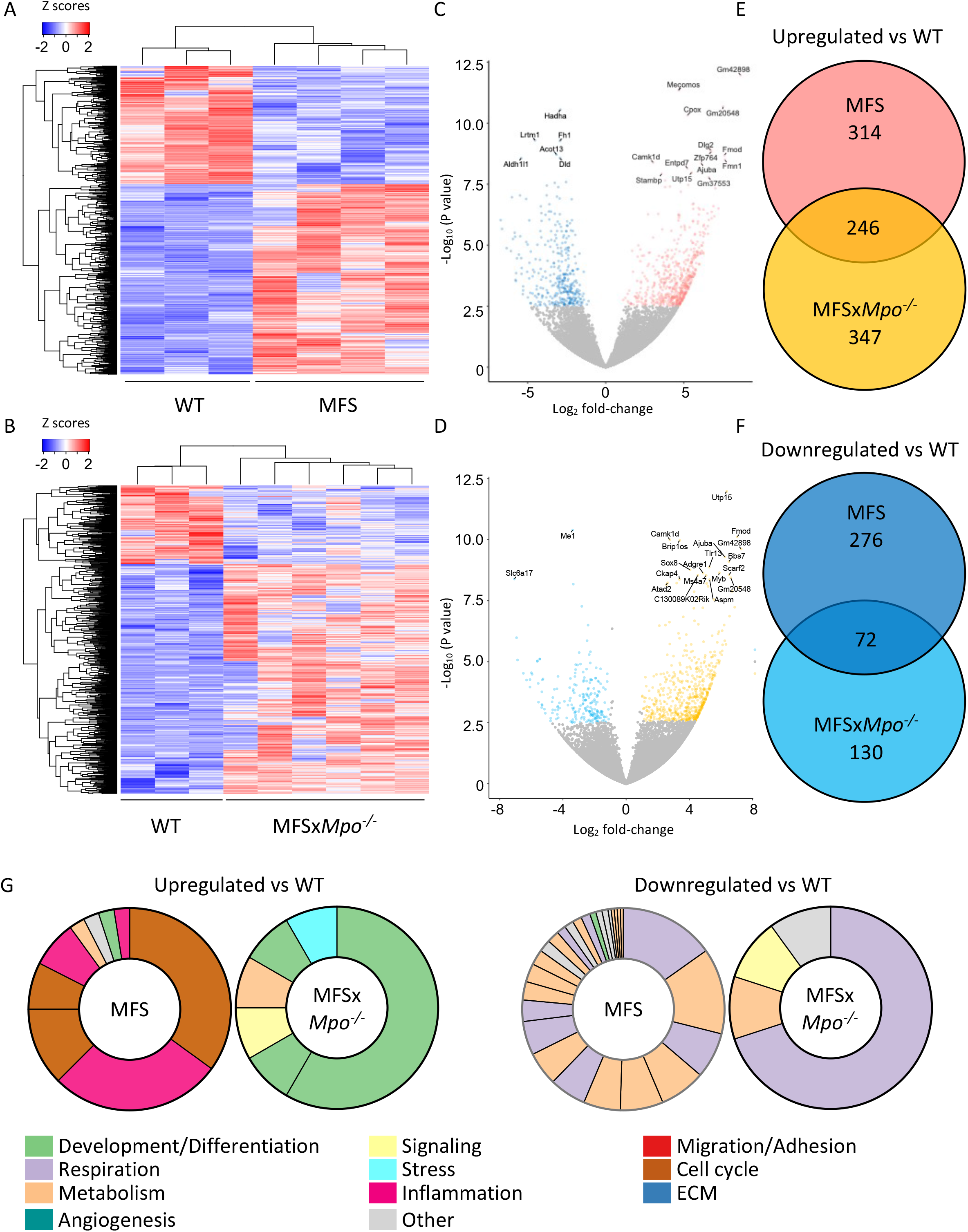
Endothelial cells show characteristic transcriptional changes in MFS. Thoracic aortic endothelial cells were isolated from the ascending aortic arch of 12-14-week-old wildtype (WT, N=3), Fbn1^C1041G/+^ (MFS, N=6), and MPO-deficient MFS (MFSx*Mpo*^-/-^, N=4) mice for bulk transcriptome analysis. Gene expression was normalized using Deseq2’s default parameters and a shifted log transformation [log10 (n+1)]. Expression was scaled across rows and the Z-scores plotted. Heatmap visualizing the significantly differentially expressed (DE) genes between (A) WT and MFS mice and (B) WT and MFSx*Mpo*^-/-^ mice. (C) Volcano plot showing the significantly up- (red) and downregulated (blue) DE genes in WT vs. MFS mice. The top 20 DE genes are labeled. (D) Volcano plot showing the significantly up- (yellow) and downregulated (light blue) DE genes between in MFSx*Mpo*^-/-^ vs. WT mice. The top 20 DE genes are labeled. (E, F) Venn diagrams indicating the individually and commonly up- (E) and downregulated (F) genes in ECs of MFS and MFSx*Mpo*^-/-^ mice in comparison to WT controls. (G) The unique differentially expressed genes were subjected to gene ontology analysis and are reported as a collapsed pie chart. The size of individual pie slices indicates the number of terms summarized within this pathway. The color indicates the function of the pathway.

Given the upregulation of inflammatory pathways of ECs in MFS mice at the transcriptional level, we next aimed to validate and further characterize the MFS-related inflammatory response in aortic ECs. Leukocytes are attracted by chemokines and need adhesion molecules including integrins for rolling, adhesion, and migration from the circulation into the inflamed tissue.^38^ Under inflammatory conditions, ECs increase expression of ICAM-1 - a member of the immunoglobulin superfamily - that is critically involved in several steps of the leukocyte migration cascade. *Von Willebrand factor*-positive aortic ECs from MFS mice showed higher immunoreactivity for ICAM-1 in comparison to WT and MFSx*Mpo*^-/-^ mice (**Fig. 5A, Sup Fig. 4**). ECs express the transmembrane protein syndecan 1, a major cell surface heparan sulfate proteoglycan within the ECM responsible for signal transduction and endothelial function.^39^ Under inflammatory conditions, the ectodomain of syndecan 1 is shedded. Accordingly, we observed a three-fold elevated plasma syndecan 1 concentration in MFS mice compared to WT and MFSx*Mpo*^--/--^ mice (**Sup Fig. 5**), further corroborating an ongoing inflammatory response of the endothelium in MFS mice. To determine the pathophysiological relevance of the observed inflammatory endothelial activation in vivo, we performed intravital microscopy of arteries within the cremaster muscle and quantified the mean endothelial adhesion time and the number of rolling leukocytes. (**Fig. 5B-D**). On average, we detected up to eight rolling leukocytes per minute in arteries of MFS mice, which was drastically reduced in MPO compound mutant MFS mice. Similarly, endothelial leukocyte adhesion was significantly increased in MFS mice compared to MFSx*Mpo*^-/- 4^ mice. We did not observe any leukocyte rolling or adhesion in WT mice. The increased rolling and adhesion coincided with a higher leukocyte content in aortic sections of MFS mice, which was reduced to WT levels in MFSx*Mpo*^-/-^ mice (**Fig. 5E**). Dihydroethidium (DHE) detects ROS, particularly superoxide. Elevated superoxide in the aorta can derive from multiple sources including leukocytes. In accordance with our findings, the DHE positive positive area was significantly increased in sections of MFS mice compared to WT controls or MFSx*Mpo*^-/-^ mice (**Fig. 5F**). Systemically, we did not detect any relevant changes for plasma concentrations of the pro-inflammatory cytokines IL-22, IL-4, IL-5, IFNγ, IL-2, IL-6, TNFa, IL-1β, IL-9, IL17-F, and IL-17A between MFS, WT, and MFSx*Mpo*^-/- 4^ mice (data not shown). Only IL-10 was increased in MFSx*Mpo^-/-^* (WT: 2.29±0.14 pg/ml, MFS: 2.37±0.05 pg/ml, MFSx*Mpo*^-/-^: 2.58±0.30 pg/ml, WT vs MFSx*Mpo*^-/-^ p= 0.01, MFS vs MFSx*Mpo*^-/-^ p=0.09).

**Figure 5.**
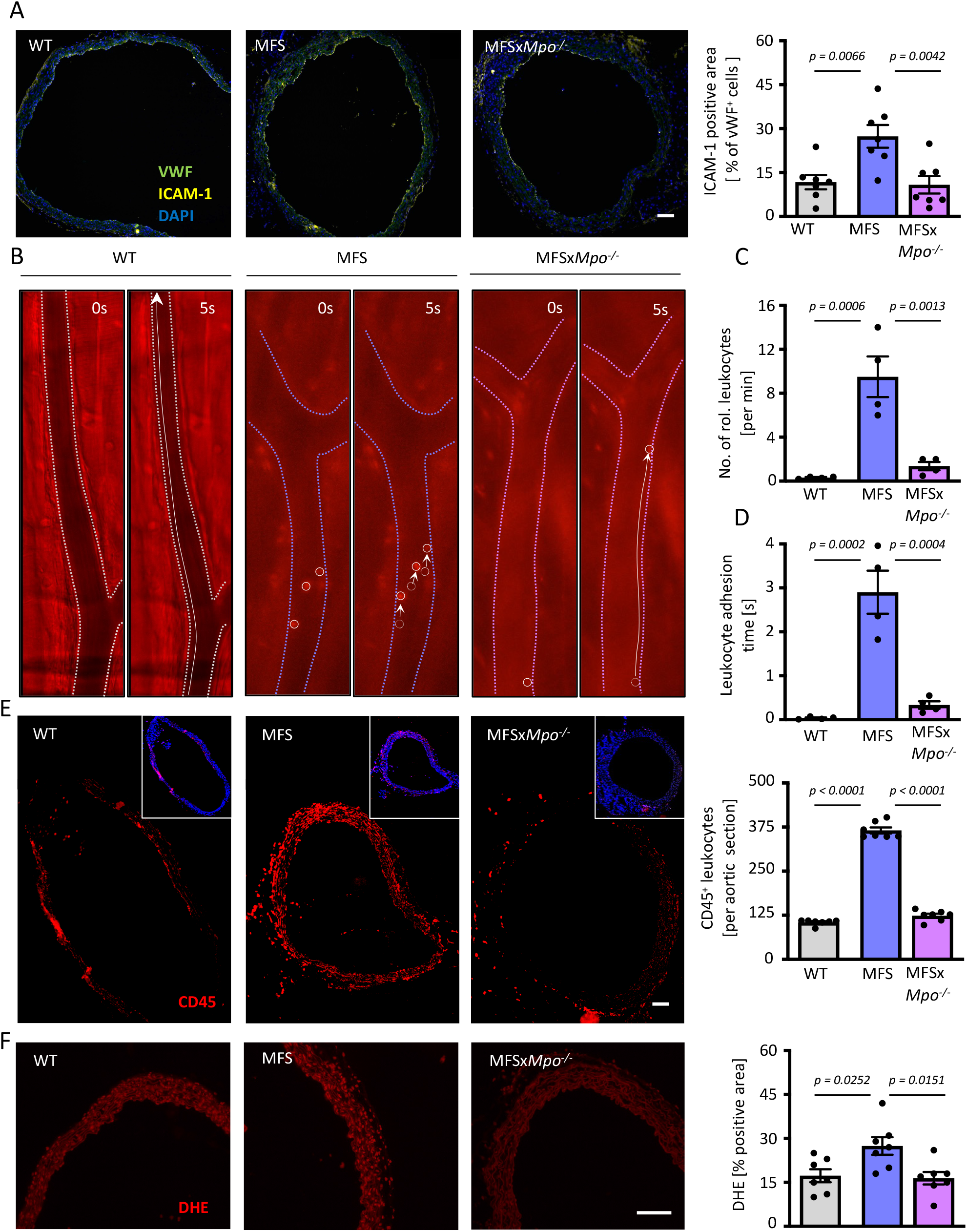
Vascular inflammation is reduced in MPO-deficient MFS mice. (A) Representative immunofluorescence images of aortic cross-sections co-stained for von-Willebrand Factor (VWF, green) and ICAM-1 (yellow). The frequency of double positive cells was quantified (N=7/group). Scale bar indicates 50 μm. (B-D) Intravital-microscopy of the cremaster muscle. Per mouse, five different fields of interest were chosen and videos recorded. Quantification of (C) rolling leukocytes per minute and (D) the adhesion of leukocytes on the endothelium in individual mice (N=4/group). (E) Representative immunofluorescence images of aortic cross-sections stained for CD45 (red) with quantification of CD45^+^ leukocytes. Overlay with DAPI is shown in the small white boxes (N=7/group). Scale bar indicates 50 μm. (F) Representative DHE staining of ascending aortic sections and quantification of aortic ROS (intracellular superoxide, red) levels as percentage of DHE positive area (red) of aortic tunica media area. Scale bar indicates 50 μm. Data are expressed as mean ± SEM. Statistical significance was determined by one-way ANOVA with Tukey’s multiple comparisons test. Wildtype (WT), Fbn1^C1041G/+^ (MFS), MPO-deficient MFS (MFSx*Mpo*^-/-^).

### Pharmacological MPO inhibition attenuates MFS progression

Finally, we tested whether pharmacological inhibition of MPO might be able to mimic the observations made for genetic ablation of MPO in MFS mice. Young MFS mice (up to 2 months of age) have a normal aortic appearance, which thereafter is progressively deteriorating and shows elastic fiber fragmentation and enlargement^40,41^, although aortic dilatation has been reported as early as 4 weeks after birth.^36^ We thus fed 6-week-old MFS mice for 6 weeks to 12 weeks of age with a diet containing the specific MPO inhibitor AZM198 or a control diet (**Fig. 6A**). AZM198 has been successfully applied and reduced disease in mouse models of nonalcoholic steatohepatitis^42^ and crescenting glomeroluonephritis^43^, where it was detectable at 1 μg/ml in plasma. Echocardiography showed that administration of the clinically available MPO inhibitor successfully attenuated thoracic aneurysm formation and aortic dilatation in MFS mice as compared to controls (**Fig. 6B-F**). On the ultrastructural level, MPO inhibition reduced elastic lamina breaks (**Fig. 6G**) while levels of inflammatory cytokines were unaffected (data not shown)

**Figure 6.**
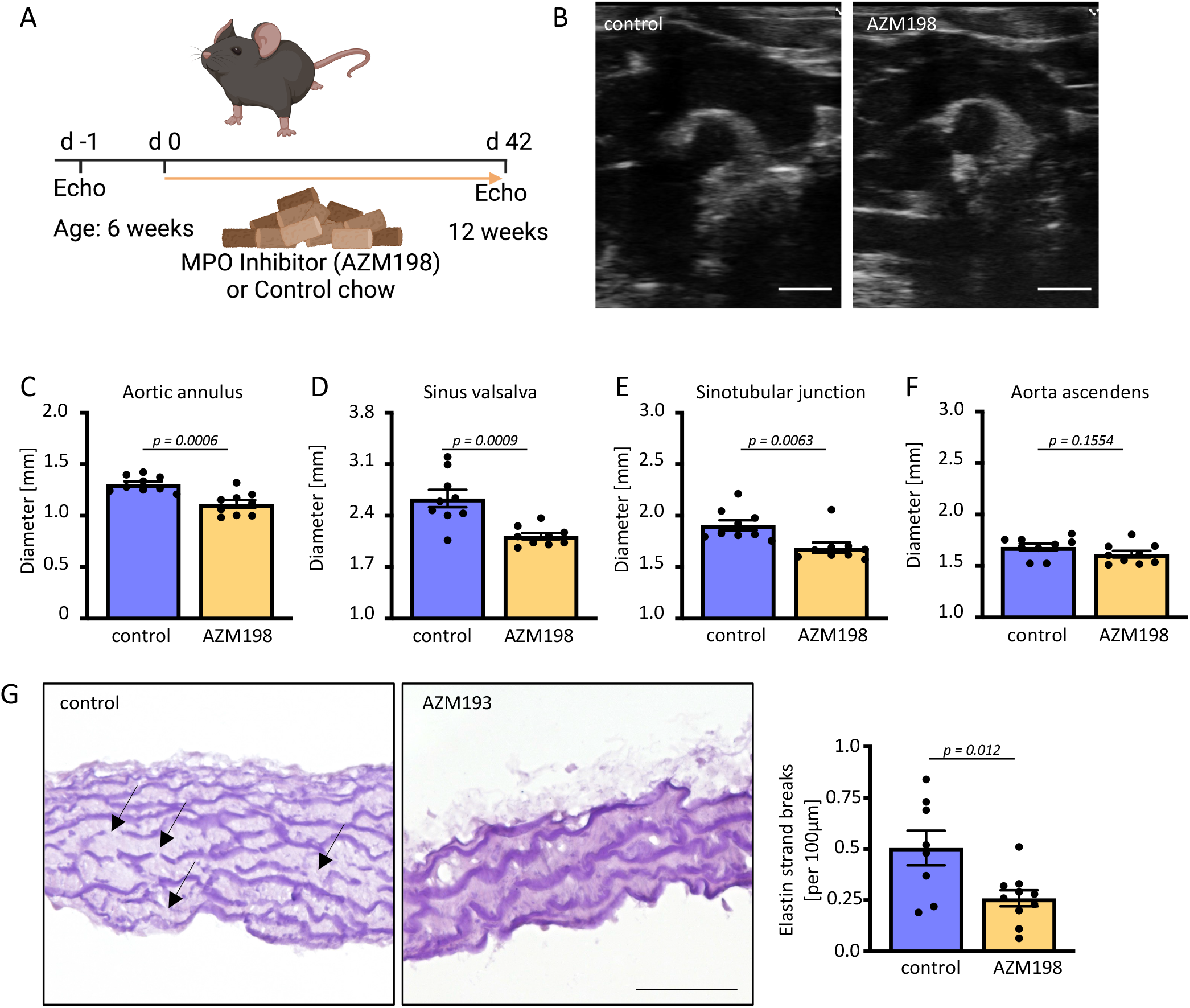
Pharmacological MPO inhibition prevents TAA formation. (A) Experimental layout. Six-week-old MFS were fed either a control diet (N=9) or a MPO inhibitor AZM198 containing diet (N=10) for six weeks. (B) Representative 2D echocardiographic image with diameter measurements at the position of the (C) aortic annulus, (D) sinus Valsalva, (E) sinotubular junction, and (F) aorta ascendens in twelve-week-old-MFS mice. Scale bar indicates 1 mm. (G) Representative images of van Gieson stained elastica in ascending aortic sections and average number of elastin breaks per 100 μm of elastic lamina (control N=9, MFS + AZM198 N=10). Scale bar indicates 50 μm. Data is presented as mean ± SEM. Statistical significance was determined by an unpaired t test.

## Discussion

We herein reveal that leucocyte-derived MPO is essentially involved in the pathogenesis of MFS-related aortic dilatation and remodeling. We could also identify pharmacological MPO inhibition as a potent treatment approach with considerable translational potential. The current data comprise the first report demonstrating the therapeutic efficacy of a targeted immunotherapy in MFS. Traditionally, adverse ECM remodeling resulting from excessive TGF-β signaling has been regarded as the major pathomechanism underlying MFS-related aortic aneurysm formation.^44^ Seminal work by Dietz and colleagues uncovered diminished TGF-β signaling by the angiotensin-II type 1 (AT1) receptor antagonist Losartan, which prevented aortic dilatation in a mouse model of MFS.^13^ Subsequent clinical trials demonstrated the capability of Angiotensin receptor blockers (ARBs) to attenuate aortic dilation in MFS patients^45^ and suggested some clinical benefit upon long-term administration.^46^ Current guidelines recommend treatment of MFS patients with ARBs as an alternative (Class I recommendation) or add-on (Class IIa recommendation) to beta blockers, the previous standard therapy.^47^ Yet, the clinical benefit of ARBs was found to be modest and failed to reach such impressive results in patients as observed in animal studies: A recent meta-analysis reported an annual change in aortic growth rate of 0.38 mm vs. 0.52 mm in patients randomized to ARBs vs. control treatment.^48^. Accumulating evidence suggests the involvement of an inflammatory response in the pathogenesis of MFS.^18,37,49–53^ However, precise understanding of causes and consequences of inflammation and particularly the role of the innate immune system in MFS is lacking, and previous findings have not yet been translated to clinically available treatment strategies specifically targeting MFS-related immune responses.

MPO is one of the most abundant proteins in neutrophils and critically involved in many cardiovascular diseases.^22^ MPO it is a potent producer of ROS, which play a crucial role in the development of TAA formation in MFS mice.^54^ Plasma MPO correlated with endothelial dysfunction in humans,^55^ which in turn correlated with aortic growth in MFS patients.^56^ We could detect increased circulating MPO levels and MPO deposition in aortas of MFS patients and mice. Moreover, we found that MPO induced an inflammatory gene transcription program in thoracic aortic endothelial cells and endothelial activation resulting in enhanced aortic leukocyte recruitment in MFS mice. In addition to increased endothelial ICAM1 expression, we observed increased plasma Syndecan 1 level in MFS mice indicative of endothelial glycocalyx shedding, which has been reported to activate leukocytes.^57^ The inflammatory EC response is in line with a recent bulk transcriptome analysis of TAA tissues from MFS patients and MFS mice, which showed a Marfan-related upregulation of pathways related to “*cell adhesion molecules*” and “*leukocyte transendothelial migration*”.^58^ In a more fine-grained in silico scRNA-seq analysis of aortic cells isolated from the same MFS mouse strain,^47^ we identified two aortic EC cluster. EC cluster 2 shared features with lymphatic endothelial cells and two previously identified aortic EC populations.^23^ The discrepancy in resolution might stem from a lower endothelial cellularity in the single cell dataset reanalyzed here. Both EC populations lost expression of several EC marker genes and acquired expression of mesenchymal cell marker genes. These are hallmark features of endothelial to mesenchymal transition (EndMT), a process predominantly driven by TGF-β signalling^31^ that contributes to various inflammatory and fibrotic disorders of the vasculature.^59^

Endothelial dysfunction and exaggerated NO signaling emerged as novel mechanisms that propagate the underlying remodeling processes in MFS.^60^ We here confirmed increased vascular aortic stiffness as redout of endothelial dysfunction in MFS mice, which is also observed in MFS patients and indicative of enhanced aortic enlargement.^61^ Our single-cell in silico analysis revealed reduced nitric oxide biosynthesis pathway activity in EC cl.1 in MFS, which has been observed previously as reduced eNOS activity^62^, and may be a trigger for the vascular smooth muscle cells to increase their iNOS expression to generate NO in MFS aortas. Indeed, the MPO/H_2_O_2_ system serves as a catalytic sink for NO, limiting its bioavailability and function and thereby upregulates inducible nitric oxide synthase, potentially further aggravating its pathological role in MFS.^63^ The excessive NO-signaling in the presence of ROS has been shown correlate with nitration of aortic peptides in mice and MFS patients.^4^. We here show that genetically ablating MPO leads to a decrease of nitration, linking increased oxidative stress and MMP activation, both together leading to increased apoptosis and elastin fragmentation in MFS aortae by MPO. ROS-dependent MMP overactivation and SMC apoptosis are established mechanisms of MFS-related TAA formation, that were recently attributed to overexpression of NAPDH oxidase and Xanthine oxidoreductase.^54,64–66^ Furthermore, ROS were shown to induce endothelial dysfunction^16^ and pro-inflammatory NF-κB signaling in MFS.^67^ Our data clearly show that MPO constitutes an important mediator of increased aortic ROS formation in MFS. Whereas MPO-derived ROS likely contributed to the observed inflammatory activation of the endothelium, non-enzymatic, electrostatic properties of MPO facilitating leukocyte recruitment could represent another important mechanism mediating aortic inflammation in MFS.^24^ Our data revealed that both, genetic MPO deficiency or pharmacological MPO inhibition sufficed to prevent MFS-related elastin fragmentation and aneurysm formation underscoring the important mechanistic role of this pathway.

In concert with previous work, our data corroborate and importantly expand the notion that the pathogenesis of MFS-related TAA formation underlies a vicious circle, in which (1) *Fbn1* deficiency gives rise to TGF-β overactivation due to insufficient ECM sequestration, (2) subsequent TGF-β-dependent EndMT and endothelial inflammatory activation promoting leukocyte adhesion and transmigration. MPO is released by recruited leukocytes or it transcytoses into the subendothelial space^20^, where (3) it accelerates endothelial dysfunction and (4) directly promotes adverse vascular remodeling by inducing oxidative stress and nitration of proteins. These processes can be mitigated by pharmacological MPO inhibition.

Our work has some limitations. Given the myriad of pro-oxidant actions of MPO, the enzyme could contribute to MFS-related aneurysm formation through additional mechanisms. We show that MPO causes nitration in the aorta, which is likely extracellular. The intracellular oxidative stress may indirectly of MPO also cause nitration of intracellular lipids and proteins and thus promote EC (and SMC) dysfunction. Nevertheless, our findings hold significant translational potential, as patients that would particularly benefit from a potential MPO inhibition could be easily identified by measuring MPO plasma levels. Pharmacological MPO inhibition targets a novel mechanism not covered by current treatment strategies and might thus provide an additive therapeutic benefit. Several oral MPO inhibitors have been developed (e.g. AZD4831 and AZD3241) and data on safety and efficacy in humans are currently under investigation, which makes their further clinical evaluation in the current disease setting feasible.^68–70^

In conclusion, we herein reveal a critical involvement of MPO in pathogenesis of MFS-related TAA formation and identify pharmacological MPO inhibition as a promising immunotherapeutic strategy in this disease. Plasma MPO could be measured prospectively to evaluate its significance as biomarker for disease prognosis. Given the advent of pharmacological MPO inhibition, the prognostic impact of the disease, and the shortcomings of current treatment approaches, therapeutic strategies targeting MPO in MFS may point towards a disease modifying pharmacological strategy in the future.

## Material and Methods (Supplement)

### Mice

Heterozygous Fbn1^C1041G/+^ mice (MFS mice) with a missense mutation in the Fbn1 gene (glycine for cysteine substitution at position 1041) on C57BL/6J background were purchased from Jackson Laboratory (#:012885) and crossbred with myeloperoxidase-deficient (*Mpo^-/-^*) mice bred in our animal facility to obtain Mpo^+/+^/Fbn1^C1041G/+^ (MFS) and *Mpo*^-/-^/Fbn1^C1041G/+^ mice (MFSx*Mpo*^-/-^). We investigated 12-14-week male mice used littermate controls. For organ collection, mice were deeply anaesthetized by inhalation of isoflurane (Isofluran-Piramal®, Piramal Critical Care, Voorschoten, The Netherlands; 5% vol/vol for induction and 2% vol/vol for maintenance of anesthesia) and subcutaneous injection of buprenorphine (TEMGESIC®, Indivior Europe Limited, Dublin, Ireland; 0.1 mg per kg body weight). The adequacy of the anesthesia was confirmed by pedal reflex testing and the animals euthanized by cardiac exsanguination.

All animal studies were approved by the local Animal Care and Use Committees [Ministry for Environment, Agriculture, Conservation and Consumer Protection of the State of North Rhine-Westphalia: State Agency for Nature, Environment and Consumer Protection (LANUV), NRW, Germany, AZ: 84-02.04.2019.A033, 84-02.04.2016.A212 and conformed to the guidelines from Directive 2010/63/EU of the European Parliament on the protection of animals used for scientific purposes.

### Echocardiography

Starting at the age of six weeks, the onset of aortic dilatation, mice were monitored by transthoracic echocardiography to 12-14 weeks of age, when the fundamental aortic remodeling has already occurred. Mice were anesthetized by isoflurane inhalation (Piramal Critical Care, 5% vol/vol for induction and 2% vol/vol for maintenance) before transthoracic echocardiography. The aorta was imaged in the B-mode aortic arch view with a Vevo 3100 ultrasound system (FUJIFILM VisualSonics, Toronto, ON, Canada) equipped with a MX550D transducer (25–55 MHz, centre transmit: 40 MHz, axial resolution: 40 μm) as previously described^65,71^. Measurements were obtained in duplicate by two blinded investigators. Aortic diameters at level of the aortic annulus, aortic root (AR), sinotubular junction and the middle ascending aorta (MAA) were measured from leading edge to leading edge at end-diastole with VevoLAB Software (FUJIFILM VisualSonics, Toronto, ON, Canada). VevoVasc (FUJIFILM VisualSonics, Toronto, ON, Canada) was used for calculation of pulse wave velocity (PWV) of the carotid artery. A B-Mode ECG-gated Kilohertz visualization (EKV) was used to define a proximal and distal position of the vessel at which a simultaneous anatomical M-mode allowed to measure the distance (d) between these points with a superior spatial and temporal resolution. Times from the ECG R wave peak to the onsets of proximal (t1) and distal carotid artery (t2) were calculated. T 1 and t2 values were averaged over 10 cardiac cycles, and the aortic arch transit time was determined as the difference of these two times (t2 – t1). Pulse wave velocity (PWV) was calculated by dividing distance (d) by the carotid artery transit time (t2 – t1) (PWV = d/(t2 – t1).

### Magnetic resonance imaging

Experiments were performed at a vertical 9.4 T Bruker AVANCEIII Wide Bore NMR spectrometer (Bruker, Ettlingen, Germany) operating at frequencies of 400.21 MHz for ^1^H using microimaging units as described previously.^72,73^ Mice were anesthetized with 1.5% isoflurane and were kept at 37°C during the measurements. For gated MRI acquisitions, the front-paws and the left hind-paw were attached to ECG electrodes (Klear-Trace) and respiration was monitored by means of a pneumatic pillow positioned at the animal’s back. Vital functions were acquired by a M1025 system (SA Instruments) and used to synchronize data acquisition with cardiac and respiratory motion. Data were acquired using a 25-mm quadrature resonator tuneable to ^1^H. To visualize the anatomy of the region of interest, ^1^H MR images were acquired using an ECG- and respiratory-gated segmented fast gradient echo cine sequence with steady-state precession (FISP). A flip angle (FA) of 15°, echo time (TE) of 1.23 ms, and a repetition time (TR) of about 6–8 ms (depending on the heart rate) were used to acquire 16 frames per heart cycle from a field of view (FOV) = 2.56 × 2.56 cm^2^, matrix = 256 × 256, 0.1 × 0.1 mm^2^ in plane resolution, 1 mm slice thickness (ST); acquisition time (TAcq) = ~2 min]. To quantify the area of the aortic annulus, the cross section either of the flowing blood (luminal diameter) and the aortic wall based on the anatomical scans was determined by manual segmentation using the region-of-interest (ROI) tool in ParaVision (Bruker). Since the aortic wall is only 100 μm in diameter, which is hardly detectable by conventional ^1^H MRI, we determined the extension of the external diameter.

### Pharmaceutical inhibitor treatment

MFS mice and wildtype littermates were either fed a standard chow diet or standard chow supplemented with the MPO inhibitor AZM198 (AstraZeneca, Stockholm, Sweden; 1,146g/kg) ad libitum. Mice were fed from the age of 6 weeks onwards. At 12 weeks of age, the thoracic aortic diameter was examined by echocardiography, analysis and tissues collected for subsequent analysis.

### Plasma collection and aortic tissue preparation

Blood was withdrawn from deeply anesthetized mice via cardiac exsanguination and collected in EDTA-coated tubes (Sarstedt, Nümbricht, Germany). Hematologic analysis was performed with a HT5 (ScilVet, Viernheim, Germany). Plasma was isolated by centrifugation (500 x g, 15 min, 4°C) and stored at −80°C until analysis. After blood withdrawal, mice were perfused with ice-cold saline and the ascending aorta was dissected. For histological analyses, the ascending aorta was embedded in Tissue-Tek optimum cutting temperature (O.C.T.) compound (Sakura Finetek, Amsterdam, The Netherlands) and frozen on dry ice. Serial sections of frozen tissue specimen with 5 μm thickness were prepared and mounted on a microscope slide. Alternatively, the adventitia was carefully removed, and the ascending aorta was snap-frozen in liquid nitrogen for further analyses.

### Histology

Selected murine aortic cryosections were histologically analyzed. Elastic fibre fragmentation was determined by Elastica-van-Gieson (EVG) staining (Elastica van Gieson staining kit, Carl Roth, Karlsruhe, Germany). Microphotographs were taken with a Keyence BZ-9000 microscope (Keyence, Osaka, Japan) at 10 x (overview of whole aortic cross-section) and 40 x magnification. Elastin breaks, defined as interruptions in the elastic fibres, were counted in four sections per mouse by a blinded investigator. The average number of elastin breaks per 100 μm elastic fibre length was calculated.

Dihydroethidium (DHE, Thermo Fisher Scientific, Waltham, MA, USA) staining of aortic cryosections was used to quantify ROS production. Tissue sections were stained with DAPI (1/1000 in 1x PBS, Thermo Fisher Scientific), incubated with DHE (5 μM, diluted in DMSO and HBSS buffer) for 30 min at 37 °C in a dark humid chamber and subsequently imaged with the Keyence BZ-9000 microscope at 40 x magnification. Dihydroethidium-positive area was planimetrically quantified by a blinded investigator using the Keyence BZ-II analyzer software.

For immunofluorescence stainings, frozen sections were thawed, fixed with 4% paraformaldehyde for 10 minutes, and blocked for 60 min with blocking solution (10 % fetal calf serum with 1 % BSA in 1 x PBS, all Thermo Scientific Fisher). Samples were then incubated with primary antibodies against CD45 (1:200, clone 30-F11, Biolegend) or ICAM1 (1:250, clone 1A29, Abcam) and van Willebrandt factor (1:100, polyclonal, # PA5-16634, Thermo Firsher Scientific) in blocking solution at 4 °C in a dark humid chamber. Subsequently, samples were incubated with corresponding secondary antibodies (VWF: Alexa Fluor 594 conjugated goat anti-rabbit, Invitrogen A-11012, 1/100; ICAM1: Alexa Fluor 488 conjugated chicken anti-mouse, Invitrogen A-21200, 1/100; CD45: Alexa Fluor 594 conjugated goat anti-rat, Invitrogen A-11007,1/100) for 60 minutes and with DAPI (1/1000, solved in 1 x PBS) for 15 min at room temperature in a dark humid chamber. Negative controls, in which the primary antibodies were omitted, were included for each staining. Images were acquired on a Keyence BZ-9000 microscope at 10 x or 40 x magnification and the CD45-positive cells or ICAM1-positve VWF-positive were quantified by a blinded investigator using the Keyence BZ-II analyzer software or Fiji/ImageJ.

### MPO staining human section

A histological section of a thoracic aortic aneurysm from a MFS patient was selected from the Biobank of the Institute of Pathology of the Institute of Pathology of the University Hospital Cologne, Germany (BioMaSOTA; Ethics-No. 13-091, approval May 2016). The study was approved by the Ethics Commission of Cologne University’s Faculty of Medicine. All experiments were performed in accordance with the guidelines of this committee and the Declaration of Helsinki. Written informed consent was obtained from all participants prior to inclusion in the study.

Samples were routinely formalin-fixed and paraffin-embedded (FFPE) according to local practice. A 4 μm thick section was cut from the FFPE tissue block and deparaffinized. Antigen retrieval was performed by boiling the sample for 20 min in a microwave oven in citrate buffer 10mM, pH 6. The samples were cooled to room temperature and washed twice in 1x TBS. Endogenous peroxidase activity was blocked by incubation with H_2_O_2_ (125μl 30% H_2_O_2_ n 50ml TBS) for 15 min at room temperature. The sample was washed twice for 5min in TBS with 0.1% Triton and unspecific binding was blocked in blocking buffer (10% Goat Serum and 1%BSA in TBS) for 1 h at room temperature. The section was incubated with the primary polyclonal anti MPO antibody (1:100, Merck Sigma-Aldrich cat.# 47915) in blocking buffer overnight at 4°C. Following two wash steps, the section was incubated with the an anti-rabbit seconday antibody conjugated with horseraddish peroxidase according to the manufacturer’s instruction (Nichirei Biosciences Inc, Tokyo, Japan) for 1 h at room temperature. MPO presence was detected by colometric conversion of the AEC substrate und nuclei were counterstained with hematoxylin. Samples were washed with tap water, mounted with Dako mounting medium (Merck, Darmstadt, Germany), and imaged with a Keyence BZ-9000 microscope at 40x resolution.

### In situ zymography

In situ zymography was performed to quantify MMP2/9 gelatinolytic activity as previously described^74^. Aortic sections were incubated with 50 μl gelatinase reaction buffer (150 mM NaCl, 5 mM CaCl2, 0.2 mM NaN3, 50 mM Tris/HCl, pH 7.6) containing 10 μg/ml DQ gelatine (Invitrogen, Carlsbad, CA, USA) for 3 h at 37 °C in a dark humid chamber. 10 mM EDTA was added as negative control to inhibit MMP2/9 activity. Images were acquired with a Keyence BZ-9000 microscope and gelatinolytic areas were planimetrically quantified by a blinded investigator using the Keyence BZ-II analyzer software.

### Gelatin zymography

Frozen aortic tissue samples were homogenized in Precellys ceramic kit tubes (1.4 mm, 0.5 ml, Bertin instruments, Montigny-le-Bretonneux, France) containing 150 μl EDTA-free Triton-X lysis buffer (20 mM Tris-HCl, 125 mM NaCl, 1% Triton X-100) with the Precellys 24 tissue homogenizer (Bertin instruments). Supernatants were collected after repetitive vortexing and centrifugation. Protein concentrations were determined using the Pierce BCA Protein Assay Kit (Thermo Fisher Scientific). 10 μg of protein was fractioned under non-reducing conditions on Novex™ 10% Zymogram Plus Gels (Thermo Fisher Scientific). Gels were developed and subsequently stained with Coomassie Blue according to the manufacturer’s instructions. Gels were visualised with a ChemiDoc XRS+ Gel Imaging System (Bio-Rad, Hercules, CA, USA) and bands were quantified by densitometry using ImageJ software. To control for equal protein loading, 10μg were loaded on a separate gel and GAPDH protein expression was determined. MMP2 activity was normalized to GAPDH protein expression. Uncropped western blot images can be found in Sup. Fig. 3.

### MPO and Syndecan 1 ELISA (mice)

Frozen tissue samples were homogenized in Precellys ceramic kit tubes (1.4 mm, 2.0 ml, Bertin Technologies, Motigny-le-Bretonneux, France) containing 200 μl ice-cold RIPA lysis buffer (150 mM NaCl, 5 mM EDTA pH 8.0, 50 mM Tris pH 8.0, 1 % NP40, 0.5 % sodium deoxycholate 0.1 % SDS) supplemented with protease and phosphatase inhibitors (Roche, Mannheim, Germany) using the Precellys 24 tissue homogenizer (Bertin Technologies). Supernatants were collected after repetitive vortexing and centrifugation. Pierce BCA Protein Assay Kit (Thermo Scientific) was used to determine protein concentration. Samples were diluted to the lowest detected concentration. Plasma and tissue MPO and Syndecan 1 levels were assessed by using the following ELISA kits according to the manufacturer’s instructions: Mouse MPO ELISA kit (HK210, Hycult Biotech, Plymouth Meeting, PA, USA), Mouse Syndecan-1 ELISA kit (ab273165, Abcam, Germany).

### Cytokine bead array

Plasma IL-22, IL-4, IL-5, IFNγ, IL-2, IL-6, TNFa, IL-1β, IL-9, IL17-F, IL-17A, and IL-10 concentrations were analyzed with Biolegend’s LEGENDplex (Biolegend, San Diego, CA, USA) according to the manufacturer’s instructions and flow cytometry (Cytek, Fremont, CA, USA).

### Plasma collection from human Marfan patients and MPO ELISA (human)

Human blood EDTA plasma samples from MFS patients were obtained at baseline visits as part of the RESVcue trial.^75^ Blood samples from age-matched, healthy controls were collected as part of the study “The influence of Myeloperoxidase-deficiency on the vascular system” (Ethics-No. 19-1580_1), which was approved November 2020 by the Ethics Commission of Cologne University’s Faculty of Medicine. All experiments were performed in accordance with the Declaration of Helsinki. Written informed consent was obtained from all participants prior to inclusion in both studies. MPO plasma levels were assessed with the Human Myeloperoxidase Quantikine ELISA Kit (DMYE00B, R&DSystems, Germany) according to the manufacturer’s instructions.

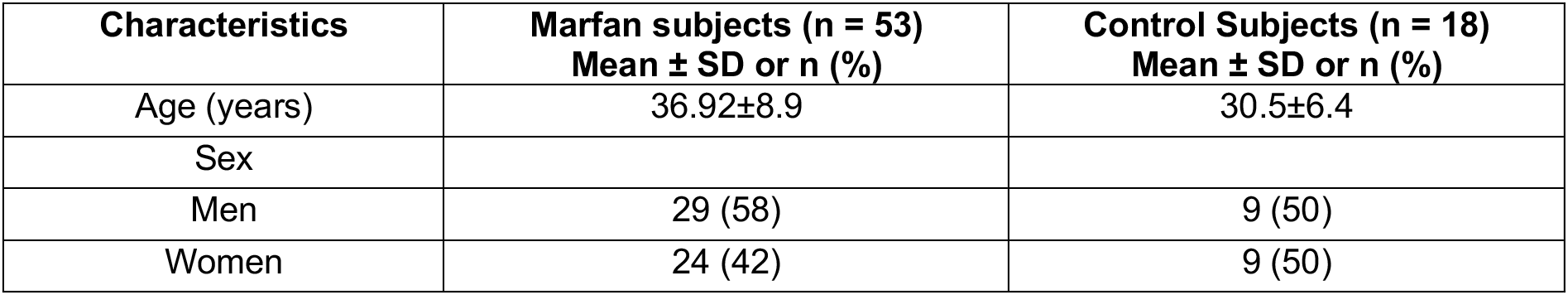

### Intravital Microscopy

Leukocyte recruitment analysis under physiological flow rates was performed by intravital microscopy of the cremaster muscle. Male, 10-14-week-old mice were anesthetized and received analgesia (100 mg/kg BW ketamine and 10 mg/kg BW xylazine i.p.). Anesthetized mice were injected i.v. via the tail vain with 100 μL 0.1% Rhodamine 6G to mark all circulating leukocytes. Mouse body temperature was maintained by placing mice on a heating plate. The fur was removed from the scrotum and the skin was incised distal from the scrotum. Subsequently, the cremaster muscle was carefully detached from external spermatic fasci and the exterior surface cleared from connective tissue. The distal end of the muscle was pinned to a custom-made silicon board. The ventral side of the cremaster was incised from distal to proximal by thermocautery. The isolated testicle was gently pushed back into the abdominal cavity. For imaging the muscle was pinned radially and flat on the silicon board and the muscle constantly kept moist with 37°C-warmed 0.9% sodium chloride. Imaging sequences of 30 sec of selected arterial vessels sized between 30μm and 50μm were taken on 5 different areas of the cremaster muscle. Intravital imaging was performed with a Leica DM6FS microscope equipped with a DFC9000 GTC camera and a 25x saline-immersion objective. Image sequences were acquired with the LasX software (Leica) and analysis of leucocyte-endothelial cell interaction was performed with Fiji/ImageJ.

### In silico single cell RNA sequencing analysis of endothelial cells in MFS

To investigate the expression profile changes of endothelial cell populations between wildtype and MFS mice, single-cell RNA-sequencing data was retrieved from GSE153534, which comprises aortic single cell transcriptomes obtained from 4-week-old and 24-week-old MFS and control mice, respectively.^30^ The paired-end FASTQ files were uploaded to the human cell atlas servers of Galaxy.^76^ Read errors were corrected and cell barcodes demultiplexed with MM3 RNA STARSolo. The raw reads of aortic single cell transcriptomes from MFS and WT mice were mapped to the annotated reference mouse (Ch39/mm39) using the STAR spliced read alignment algorithm.^77^ Next, error correction and deduplication of unique molecular identifiers was performed. The by STARsolo obtained output files were subjected for a quality control analysis and single cell transcriptome clustering and visualization in a uniform manifold approximation and projection space (UMAP) with Seurat.^76^ Cells containing less than 500 or more than 15000 transcripts were excluded. We next merged the different single cell data with the FindIntegrationAnchor function in Seurat. Cells were clustered based on the first thirty PCA dimensions and the cluster resolution was set to 0.8 in the FindCluster function in Seurat. The top cluster-identifying genes were identified for cluster calling.^78^ We next performed differential gene expression analysis and subjected the up- and downregulated genes filtered for significance p<0.05 and a log-fold change (>0.5 or <-0.5) to pathway analysis with ClueGO.^79^

### RNA isolation of murine ECs from the ascending aortic arch

Aortas from twelve-week-old male WT, MFS, and MFSx *Mpo*^-/-^ mice were dissected. The ascending aortic arch spanning from the aortic valve to the area between the right and left carotid arteries was collected. Aortic rings were cut open. ECs and adventitial cells were isolated using an ice-cold metal rod (6 mm diameter) according to the “modified Häutchen method” described by Simmons et al.^80^. RNA from ECs and adventitial cells was isolated using the RNeasy micro kit according to the manufacturer’s instructions (Qiagen, Hilden, Germany). To assess RNA quality the RNA integrity number (RIN) was determined by a 2100 Bioanalyzer (Agilent Technologies, Santa Clara, CA, USA). Only samples with a RIN above 5.6 were processed further.

### Bulk transcriptome analysis of ECs

For library preparation, the Trio RNA-Seq Library Preparation kit (TECAN, Männedorf, Switzerland) was used. Five PCR cycles were applied for library amplification and libraries with an average fragment size of 326 bp were sequenced on a NextSeq 500/ NextSeq 2000 in paired-end mode (75 bp/ 65 bp, Illumina) at the GeneCore sequencing service of the EMBL (Heidelberg, Germany). For bioinformatic analysis, we used the Galaxy platform^81^. RNA sequencing reads were mapped using RNA STAR ^77^ followed by counting reads per gene by using featureCounts ^82^. As an additional quality control step the purity of ECs in the respective sample was determined by analyzing expression levels of the classical EC marker genes *Pecam1, vWF* and *Cdh5*. The normalized counts of these 3 marker genes were added up for each sample and compared with EC marker expression in adventitial samples from 4 control animals. Only samples with EC marker expression of >2-fold of the mean EC marker expression in adventitial samples were included in the analysis. Thus, for bioinformatic analysis the following sample numbers were used: WT N=3, MFS N=4, MFSx *Mpo*^-/-^ N=6. In these samples, differentially expressed genes were identified by DESeq2 ^83^ For data visualization, normalization, and cluster analysis heatmap2 and Volcano plots Galaxy^81^ was used. Gene ontology analysis of the up and downregulated genes filtered for significance p<0.05 and a log-fold change (>0.5 or <-0.5) was performed with ClueGO using the GO-term database with the sub-ontologies “biological processes, cellular-component and molecular function”.^79^

### Statistical analysis

Data are presented as mean ± SEM. Shapiro–Wilk test and Brown–Forsythe test were utilized to test for normal distribution and equality of variances, respectively. Differences between three groups were evaluated using one-way or two-way repeated measures analysis of variance (ANOVA) with post-hoc Tukey’s test if data were normally distributed and variances were equal. Kruskal–Wallis test with Dunn’s post-hoc test was performed if data were not normally distributed. For comparison of two data sets consisting of normally distributed data, the unpaired t-test was utilized. A P-value <0.05 was considered statistically significant. All statistical analyses were performed using GraphPad Prism 8.4.0 (GraphPad Software, San Diego, CA, USA).

## Supporting information

All supplemental figures

## Acknowledgments

The authors would like to thank Christina Vosen, Sharon Weingarten, Katharina Tinaz, Nadja Klein, and Simon Grimm for excellent technical support. We further would like to thank Erik Michaelsson and Sepideh Hagvall from Astra Zeneca for providing the MPO Inhibitor AZM198.

**Sup Fig. 1**

Frequency of circulating neutrophils in Wildtype (WT, N=4), Fbn1^C1041G/+^ (MFS, N=5), MPO-deficient MFS (MFSx*Mpo*^-/-^ N=4). Data is presented as mean ± SEM. Statistical significance was determined by ordinary one-way ANOVA followed by Tukey’s multiple comparison test.

**Sup Fig. 2**

In gel zymography for MMP2/MMP9 detection.

**Sup Fig. 3**

(A,B) In silico analysis of aortic single cell transcriptomes obtained from WT and MFS mice GSE153534. Two endothelial cell cluster were identified and marker genes for all major populations are reported. Venn diagrams indicate the number of individually and jointly (C) upregulated and (D) downregulated genes in EC cluster 1 and 2 in MFS vs WT mice. Gene ontology analysis of genes exclusively (E) increased or decreased (F) in EC clusters 1 and 2 in MFS vs WT. Wildtype (WT), Fbn1^C1041G/+^ (MFS)

**Sup Fig 4.**

Representative photomicrographs of VWF (green), ICAM1 (yellow), and nuclei (blue) immunofluorescence stainings in the aortas of Wildtype (WT), Fbn1^C1041G/+^ (MFS), and MPO-deficient MFS mice. A negative control lacking incubation with the primary antibody against ICAM1 is provided.

**Sup Fig. 5**

Plasma Syndecan 1 levels were measured by ELISA (WT N=16, MFS N=8, MFSx*Mpo*^-/-^, N=8). Data are expressed as mean ± SEM. Statistical significance was determined by Kruskal-Wallis-test with Dunn’s multiple comparisons test. Wildtype (WT), Fbn1C1041G/+ (MFS), MPO-deficient MFS (MFSx*Mpo*^-/-^).

## References

1. Ramachandra, C.J., et al. Molecular pathogenesis of Marfan syndrome. Int J Cardiol 187, 585–591 (2015).

2. Dietz, H.C., et al. Marfan syndrome caused by a recurrent de novo missense mutation in the fibrillin gene. Nature 352, 337–339 (1991).

3. Wanga, S., et al. Aortic microcalcification is associated with elastin fragmentation in Marfan syndrome. J Pathol 243, 294–306 (2017).

4. de la Fuente-Alonso, A., et al. Aortic disease in Marfan syndrome is caused by overactivation of sGC-PRKG signaling by NO. Nat Commun 12, 2628 (2021).

5. Booms, P., et al. RGD-containing fibrillin-1 fragments upregulate matrix metalloproteinase expression in cell culture: a potential factor in the pathogenesis of the Marfan syndrome. Hum Genet 116, 51–61 (2005).

6. Chung, A.W., et al. Loss of elastic fiber integrity and reduction of vascular smooth muscle contraction resulting from the upregulated activities of matrix metalloproteinase-2 and −9 in the thoracic aortic aneurysm in Marfan syndrome. Circ Res 101, 512–522 (2007).

7. Segura, A.M., et al. Immunohistochemistry of matrix metalloproteinases and their inhibitors in thoracic aortic aneurysms and aortic valves of patients with Marfan’s syndrome. Circulation 98, II331–337; discussion II337-338 (1998).

8. Davis, M.R. & Summers, K.M. Structure and function of the mammalian fibrillin gene family: implications for human connective tissue diseases. Mol Genet Metab 107, 635–647 (2012).

9. Wang, Y., et al. TGF-beta activity protects against inflammatory aortic aneurysm progression and complications in angiotensin II-infused mice. J Clin Invest 120, 422–432 (2010).

10. Cook, J.R., et al. Dimorphic effects of transforming growth factor-beta signaling during aortic aneurysm progression in mice suggest a combinatorial therapy for Marfan syndrome. Arterioscler Thromb Vasc Biol 35, 911–917 (2015).

11. Wei, H., et al. Aortopathy in a Mouse Model of Marfan Syndrome Is Not Mediated by Altered Transforming Growth Factor beta Signaling. J Am Heart Assoc 6(2017).

12. Holm, T.M., et al. Noncanonical TGFbeta signaling contributes to aortic aneurysm progression in Marfan syndrome mice. Science 332, 358–361 (2011).

13. Hara, H., et al. Inhibition of transforming growth factor-beta signaling in myeloid cells ameliorates aortic aneurysmal formation in Marfan syndrome. PLoS One 15, e0239908 (2020).

14. Fiorillo, C., et al. Evidence for oxidative stress in plasma of patients with Marfan syndrome. Int J Cardiol 145, 544–546 (2010).

15. Gomez, D., et al. Epigenetic control of vascular smooth muscle cells in Marfan and non-Marfan thoracic aortic aneurysms. Cardiovasc Res 89, 446–456 (2011).

16. Jimenez-Altayo, F., et al. Redox stress in Marfan syndrome: Dissecting the role of the NADPH oxidase NOX4 in aortic aneurysm. Free Radic Biol Med 118, 44–58 (2018).

17. Guo, G., et al. Induction of macrophage chemotaxis by aortic extracts of the mgR Marfan mouse model and a GxxPG-containing fibrillin-1 fragment. Circulation 114, 1855–1862 (2006).

18. Franken, R., et al. No beneficial effect of general and specific anti-inflammatory therapies on aortic dilatation in Marfan mice. PLoS One 9, e107221 (2014).

19. Astern, J.M., et al. Myeloperoxidase interacts with endothelial cell-surface cytokeratin 1 and modulates bradykinin production by the plasma Kallikrein-Kinin system. Am J Pathol 171, 349–360 (2007).

20. Baldus, S., et al. Endothelial transcytosis of myeloperoxidase confers specificity to vascular ECM proteins as targets of tyrosine nitration. J Clin Invest 108, 1759–1770 (2001).

21. Baldus, S., et al. Spatial mapping of pulmonary and vascular nitrotyrosine reveals the pivotal role of myeloperoxidase as a catalyst for tyrosine nitration in inflammatory diseases. Free Radic Biol Med 33, 1010 (2002).

22. Kargapolova, Y., et al. The Enzymatic and Non-Enzymatic Function of Myeloperoxidase (MPO) in Inflammatory Communication. Antioxidants (Basel) 10(2021).

23. Manchanda, K., et al. MPO (Myeloperoxidase) Reduces Endothelial Glycocalyx Thickness Dependent on Its Cationic Charge. Arterioscler Thromb Vasc Biol 38, 1859–1867 (2018).

24. Klinke, A., et al. Myeloperoxidase attracts neutrophils by physical forces. Blood 117, 1350–1358 (2011).

25. Senders, M.L. & Mulder, W.J.M. Targeting myeloperoxidase in inflammatory atherosclerosis. Eur Heart J 39, 3311–3313 (2018).

26. Cheng, D., et al. Inhibition of MPO (Myeloperoxidase) Attenuates Endothelial Dysfunction in Mouse Models of Vascular Inflammation and Atherosclerosis. Arterioscler Thromb Vasc Biol 39, 1448–1457 (2019).

27. Gaut, J.P., et al. Myeloperoxidase produces nitrating oxidants in vivo. J Clin Invest 109, 1311–1319 (2002).

28. Fu, X., Kassim, S.Y., Parks, W.C. & Heinecke, J.W. Hypochlorous acid generated by myeloperoxidase modifies adjacent tryptophan and glycine residues in the catalytic domain of matrix metalloproteinase-7 (matrilysin): an oxidative mechanism for restraining proteolytic activity during inflammation. J Biol Chem 278, 28403–28409 (2003).

29. Shen, M., et al. Divergent roles of matrix metalloproteinase 2 in pathogenesis of thoracic aortic aneurysm. Arterioscler Thromb Vasc Biol 35, 888–898 (2015).

30. Pedroza, A.J., et al. Single-Cell Transcriptomic Profiling of Vascular Smooth Muscle Cell Phenotype Modulation in Marfan Syndrome Aortic Aneurysm. Arterioscler Thromb Vasc Biol 40, 2195–2211 (2020).

31. Kalluri, A.S., et al. Single-Cell Analysis of the Normal Mouse Aorta Reveals Functionally Distinct Endothelial Cell Populations. Circulation 140, 147–163 (2019).

32. Inoue, M., et al. Endothelial cell-selective adhesion molecule modulates atherosclerosis through plaque angiogenesis and monocyte-endothelial interaction. Microvasc Res 80, 179–187 (2010).

33. Korhonen, E.A., et al. Tie1 controls angiopoietin function in vascular remodeling and inflammation. J Clin Invest 126, 3495–3510 (2016).

34. Prevo, R., Banerji, S., Ferguson, D.J., Clasper, S. & Jackson, D.G. Mouse LYVE-1 is an endocytic receptor for hyaluronan in lymphatic endothelium. J Biol Chem 276, 19420–19430 (2001).

35. Weber, M., et al. Interstitial dendritic cell guidance by haptotactic chemokine gradients. Science 339, 328–332 (2013).

36. Oller, J., et al. Extracellular Tuning of Mitochondrial Respiration Leads to Aortic Aneurysm. Circulation 143, 2091–2109 (2021).

37. Verhagen, J.M.A., et al. Multi-Omics Profiling in Marfan Syndrome: Further Insights into the Molecular Mechanisms Involved in Aortic Disease. Int J Mol Sci 23(2021).

38. Ley, K., Laudanna, C., Cybulsky, M.I. & Nourshargh, S. Getting to the site of inflammation: the leukocyte adhesion cascade updated. Nat Rev Immunol 7, 678–689 (2007).

39. Afratis, N.A., et al. Syndecans - key regulators of cell signaling and biological functions. FEBS J 284, 27–41 (2017).

40. Judge, D.P., et al. Evidence for a critical contribution of haploinsufficiency in the complex pathogenesis of Marfan syndrome. J Clin Invest 114, 172–181 (2004).

41. Hibender, S., et al. Resveratrol Inhibits Aortic Root Dilatation in the Fbn1C1039G/+ Marfan Mouse Model. Arterioscler Thromb Vasc Biol 36, 1618–1626 (2016).

42. Koop, A.C., et al. Therapeutic Targeting of Myeloperoxidase Attenuates NASH in Mice. Hepatol Commun 4, 1441–1458 (2020).

43. Antonelou, M., et al. Therapeutic Myeloperoxidase Inhibition Attenuates Neutrophil Activation, ANCA-Mediated Endothelial Damage, and Crescentic GN. J Am Soc Nephrol 31, 350–364 (2020).

44. Habashi, J.P., et al. Losartan, an AT1 antagonist, prevents aortic aneurysm in a mouse model of Marfan syndrome. Science 312, 117–121 (2006).

45. Lacro, R.V., et al. Atenolol versus losartan in children and young adults with Marfan’s syndrome. N Engl J Med 371, 2061–2071 (2014).

46. van Andel, M.M., et al. Long-term clinical outcomes of losartan in patients with Marfan syndrome: follow-up of the multicentre randomized controlled COMPARE trial. Eur Heart J 41, 4181–4187 (2020).

47. Sellers, S.L., et al. Inhibition of Marfan Syndrome Aortic Root Dilation by Losartan: Role of Angiotensin II Receptor Type 1-Independent Activation of Endothelial Function. Am J Pathol 188, 574–585 (2018).

48. Pitcher, A., et al. Angiotensin receptor blockers and beta blockers in Marfan syndrome: an individual patient data meta-analysis of randomised trials. Lancet 400, 822–831 (2022).

49. Lim, W.W., et al. Inhibition of IL11 Signaling Reduces Aortic Pathology in Murine Marfan Syndrome. Circ Res 130, 728–740 (2022).

50. Guido, M.C., et al. Treatment With Methotrexate Associated With Lipid Core Nanoparticles Prevents Aortic Dilation in a Murine Model of Marfan Syndrome. Front Cardiovasc Med 9, 893774 (2022).

51. Zhang, R.M., et al. Fibrillin-1-regulated miR-122 has a critical role in thoracic aortic aneurysm formation. Cell Mol Life Sci 79, 314 (2022).

52. Ju, X., et al. IL-6 regulates extracellular matrix remodeling associated with aortic dilation in a fibrillin-1 hypomorphic mgR/mgR mouse model of severe Marfan syndrome. J Am Heart Assoc 3, e000476 (2014).

53. Radonic, T., et al. Inflammation aggravates disease severity in Marfan syndrome patients. PLoS One 7, e32963 (2012).

54. Emrich, F., et al. Anatomically specific reactive oxygen species production participates in Marfan syndrome aneurysm formation. J Cell Mol Med 23, 7000–7009 (2019).

55. Mollenhauer, M., et al. Myeloperoxidase Mediates Postischemic Arrhythmogenic Ventricular Remodeling. Circ Res 121, 56–70 (2017).

56. Takata, M., et al. Impairment of flow-mediated dilation correlates with aortic dilation in patients with Marfan syndrome. Heart Vessels 29, 478–485 (2014).

57. Vita, J.A., et al. Serum myeloperoxidase levels independently predict endothelial dysfunction in humans. Circulation 110, 1134–1139 (2004).

58. Quarto, N., Li, S., Renda, A. & Longaker, M.T. Exogenous activation of BMP-2 signaling overcomes TGFbeta-mediated inhibition of osteogenesis in Marfan embryonic stem cells and Marfan patient-specific induced pluripotent stem cells. Stem Cells 30, 2709–2719 (2012).

59. Alvandi, Z. & Bischoff, J. Endothelial-Mesenchymal Transition in Cardiovascular Disease. Arterioscler Thromb Vasc Biol 41, 2357–2369 (2021).

60. Oller, J., et al. Nitric oxide mediates aortic disease in mice deficient in the metalloprotease Adamts1 and in a mouse model of Marfan syndrome. Nat Med 23, 200–212 (2017).

61. Selamet Tierney, E.S., et al. Influence of Aortic Stiffness on Aortic-Root Growth Rate and Outcome in Patients With the Marfan Syndrome. Am J Cardiol 121, 1094–1101 (2018).

62. Chung, A.W., et al. Endothelial dysfunction and compromised eNOS/Akt signaling in the thoracic aorta during the progression of Marfan syndrome. Br J Pharmacol 150, 1075–1083 (2007).

63. Galijasevic, S., Saed, G.M., Diamond, M.P. & Abu-Soud, H.M. Myeloperoxidase up-regulates the catalytic activity of inducible nitric oxide synthase by preventing nitric oxide feedback inhibition. Proc Natl Acad Sci U S A 100, 14766–14771 (2003).

64. Emrich, F.C., et al. Enhanced caspase activity contributes to aortic wall remodeling and early aneurysm development in a murine model of Marfan syndrome. Arterioscler Thromb Vasc Biol 35, 146–154 (2015).

65. Nettersheim, F.S., et al. Nitro-oleic acid (NO2-OA) reduces thoracic aortic aneurysm progression in a mouse model of Marfan syndrome. Cardiovasc Res (2021).

66. Rodriguez-Rovira, I., et al. Allopurinol blocks aortic aneurysm in a mouse model of Marfan syndrome via reducing aortic oxidative stress. Free Radic Biol Med (2022).

67. You, W., et al. TGF-beta mediates aortic smooth muscle cell senescence in Marfan syndrome. Aging (Albany NY) 11, 3574–3584 (2019).

68. Nelander, K., et al. Early Clinical Experience With AZD4831, A Novel Myeloperoxidase Inhibitor, Developed for Patients With Heart Failure With Preserved Ejection Fraction. Clin Transl Sci 14, 812–819 (2021).

69. Jucaite, A., et al. Effect of the myeloperoxidase inhibitor AZD3241 on microglia: a PET study in Parkinson’s disease. Brain 138, 2687–2700 (2015).

70. Gan, L.M., et al. Safety, tolerability, pharmacokinetics and effect on serum uric acid of the myeloperoxidase inhibitor AZD4831 in a randomized, placebo-controlled, phase I study in healthy volunteers. Br J Clin Pharmacol 85, 762–770 (2019).

71. Lee, L., et al. Aortic and Cardiac Structure and Function Using High-Resolution Echocardiography and Optical Coherence Tomography in a Mouse Model of Marfan Syndrome. PLoS One 11, e0164778 (2016).

72. Flogel, U., et al. In vivo monitoring of inflammation after cardiac and cerebral ischemia by fluorine magnetic resonance imaging. Circulation 118, 140–148 (2008).

73. Temme, S., et al. Beyond Vessel Diameters: Non-invasive Monitoring of Flow Patterns and Immune Cell Recruitment in Murine Abdominal Aortic Disorders by Multiparametric MRI. Front Cardiovasc Med 8, 750251 (2021).

74. Gkantidis, N., Katsaros, C. & Chiquet, M. Detection of gelatinolytic activity in developing basement membranes of the mouse embryo head by combining sensitive in situ zymography with immunolabeling. Histochem Cell Biol 138, 557–571 (2012).

75. Andel, M.v., et al. The Effect of Resveratrol on Aortic Function in Patients with Marfan Syndrome – Rationale and Design of the Resvcue Marfan Trial. (Research Square, 2021).

76. Hao, Y., et al. Integrated analysis of multimodal single-cell data. Cell 184, 3573–3587 e3529 (2021).

77. Dobin, A., et al. STAR: ultrafast universal RNA-seq aligner. Bioinformatics 29, 15–21 (2013).

78. Tekman, M., et al. A single-cell RNA-sequencing training and analysis suite using the Galaxy framework. Gigascience 9(2020).

79. Bindea, G., et al. ClueGO: a Cytoscape plug-in to decipher functionally grouped gene ontology and pathway annotation networks. Bioinformatics 25, 1091–1093 (2009).

80. Simmons, C.A., Zilberberg, J. & Davies, P.F. A rapid, reliable method to isolate high quality endothelial RNA from small spatially-defined locations. Ann Biomed Eng 32, 1453–1459 (2004).

81. Afgan, E., et al. The Galaxy platform for accessible, reproducible and collaborative biomedical analyses: 2018 update. Nucleic Acids Res 46, W537–W544 (2018).

82. Liao, Y., Smyth, G.K. & Shi, W. featureCounts: an efficient general purpose program for assigning sequence reads to genomic features. Bioinformatics 30, 923–930 (2014).

83. Love, M.I., Huber, W. & Anders, S. Moderated estimation of fold change and dispersion for RNA-seq data with DESeq2. Genome Biol 15, 550 (2014).

