## Supplementary material for "Inhibition of myeloperoxidase attenuates thoracic aortic aneurysm formation in Marfan disease": All supplemental figures

### Supplementary Figure 1

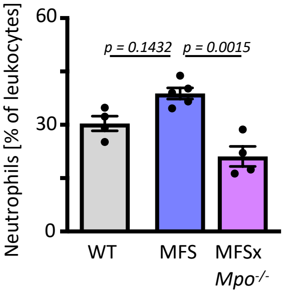

### Supplementary Figure 2

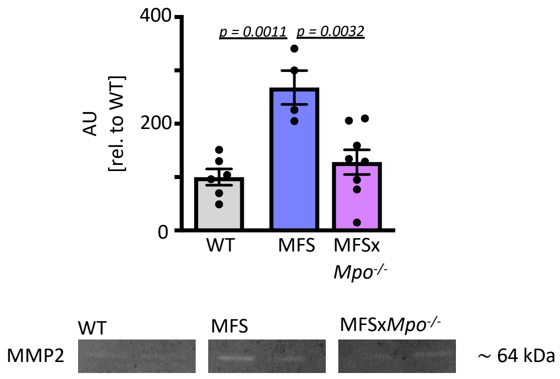

### Supplementary Figure 3

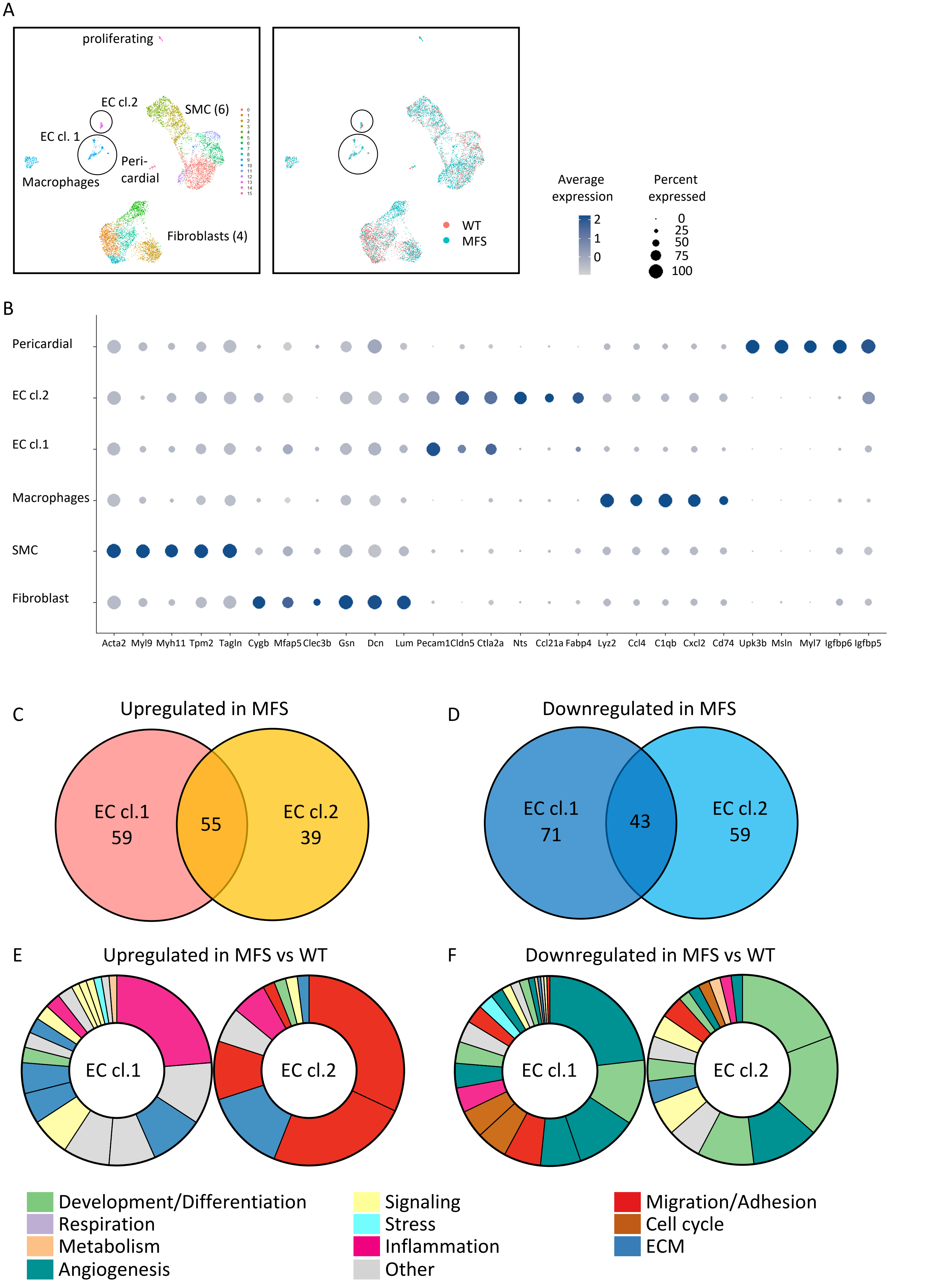

### Supplementary Figure 4

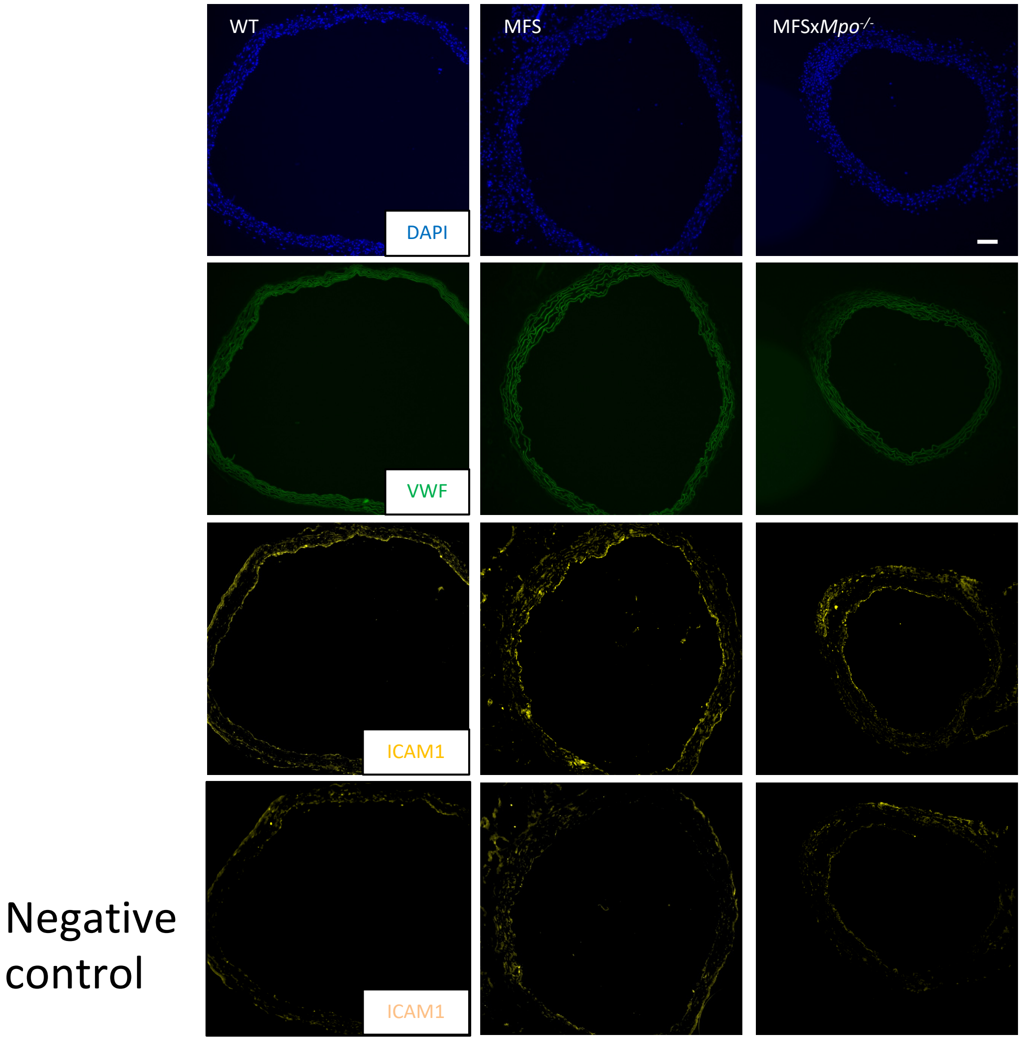

### Supplementary Figure 5

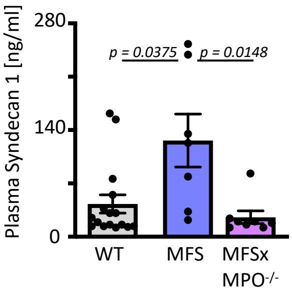
